# Functional analysis of novel microneme proteins from *Plasmodium vivax* blood stages identifies vaccine candidates

**DOI:** 10.64898/2026.08.07.743484

**Authors:** Arunaditya Deshmukh, Francisco Martinez, Pailene S. Lim, Lionel Brice Feufack-Donfack, Florent Dingli, Kaitlin Pekin, Baura Tat, Benson Kiniboro, D. Herbert Opi, Yee-Ling Lau, Mun Yik Fong, Eun-Taek Han, James G. Beeson, Jetsumon Sattabongkot, Ivo Mueller, Loew Damarys, Jean Popovici, Rhea J. Longley, Chetan E. Chitnis

## Abstract

Host cell invasion by malaria parasites requires specific molecular interactions with host receptors. *Plasmodium vivax* merozoite invasion of reticulocytes is mediated by *P. vivax* Duffy binding protein (PvDBP) and its homolog, *P. vivax* erythrocyte binding protein (PvEBP). Here, we identify and characterize two novel *P. vivax* merozoite proteins, PvMP45 and PvMP36, which co- localize with PvDBP and PvEBP in the micronemes and bind reticulocyte receptors. PvMP45 and PvMP36 share high sequence identity with their *P. knowlesi* homologs, PkMP45 and PkMP36, which form a complex with other invasion related proteins. Field studies reveal that naturally acquired antibodies against PvMP36, PvEBP and PvDBP are associated with protection against clinical *P. vivax* malaria. We demonstrate that naturally acquired antibodies to PvEBP bind Fcy receptors and likely mediate protection by enabling opsonic phagocytosis. In addition, we show that combining antibodies against PvDBP and PvMP36 results in an additive invasion inhibitory effect against *P. vivax* blood stages. These results suggest that combining PvDBP, PvEBP and PvMP36 in a multivalent blood stage vaccine could elicit diverse immune mechanisms against *P. vivax* to achieve high efficacy.

**Importance:** All the clinical symptoms of malaria are attributed to the blood stage of malaria parasites during which merozoites invade and multiply within red blood cells. A clear understanding of the host-parasite interactions that enable invasion can open paths for development of novel methods to block parasite growth and prevent malaria. Here, we identify and characterize two novel invasion related proteins from *P. vivax* merozoites that form an invasion complex and bind host RBC receptors. We demonstrate that antibodies targeting these proteins can block RBC invasion by *P. vivax* and naturally acquired antibodies that develop following *P. vivax* infection against one of these proteins are associated with protection against *P. vivax* malaria. These studies not only expand our understanding of the molecular mechanisms that enable host cell invasion by *P. vivax* but open new avenues for development of vaccines to protect against *P. vivax* malaria.

## INTRODUCTION

According to the World Health Organization, there were 282 million malaria cases worldwide in 2024, which resulted in 610,000 deaths (1). More than 90% of malaria cases were caused by *Plasmodium falciparum* infections occurring in sub-Saharan Africa with majority of deaths reported in young children less than 5 years old (1)*. Plasmodium vivax*, the other important *Plasmodium* species for human malaria, is geographically more widespread and accounts for a significant proportion of malaria cases in the horn of Africa, South and South-East Asia, the Pacific islands and South and Central America (1). *P. vivax* infections were thought to be restricted to Duffy positive individuals who express the Duffy antigen receptor (DARC) on reticulocytes (2). However, low density, asymptomatic *P. vivax* infections have been frequently reported in Duffy- negative populations from multiple locations across sub-Saharan Africa in recent years (3–6). *P. vivax* blood stage infections in Duffy-negative individuals are attributed to the transient expression of DARC in Duffy negative erythroblasts during terminal erythroid differentiation, which likely enables *P. vivax* invasion by the Duffy pathway (7–9).

Current malaria control tools are less effective at controlling *P. vivax* compared to *P. falciparum* due to its unique biology (10). This includes the ability of *P. vivax* to form dormant hypnozoites in the liver that can emerge to cause blood stage infection weeks, months or even years later. The lack of methods to detect individuals carrying hypnozoites makes *P. vivax* control difficult. Additionally, *P. vivax* produces infectious gametocytes early in blood stage infection compared to *P. falciparum*. As a result, by the time a *P. vivax*-infected patient seeks treatment, transmission may have already occurred, making early diagnosis and treatment strategies less effective.

First generation vaccines for *P. falciparum* malaria, RTS, S and R21, based on the circumsporozoite protein (PfCSP), are currently being rolled out in Africa for delivery to children (11, 12). A vaccine candidate, Rv21, based on PvCSP that is similar in design to R21, has demonstrated promising results in pre-clinical studies but has not yet been tested in clinical trials (13). The leading *P. vivax* blood stage vaccine candidate is based on the receptor-binding domain, region II, of the *P. vivax* Duffy binding protein (PvDBPII). Immunization with recombinant PvDBPII protein formulated with the adjuvant Matrix M (PvDBPII-Matrix M) in a Phase II human blood stage challenge trial reduced the parasite multiplication rate *in vivo* by 53% (14–15). This study provides the first evidence demonstrating the potential of a vaccine based on PvDBPII to control *P. vivax* blood stage infection in humans. However, there is a need to achieve higher levels of blood stage growth inhibition. This could be achieved by combining PvDBPII with other key *P. vivax* blood stage antigens that play a role in reticulocyte invasion. In addition to PvDBP, *P. vivax* merozoites express a homolog that is referred to as *P. vivax* erythrocyte binding protein (PvEBP) (16–18). PvEBP has been shown to bind complement receptor 1 (CR1) on RBCs (19) but its role in invasion is not clear. The binding domain of PvEBP also lies in the conserved N-terminal cysteine-rich domain, region II (PvEBPII), that shares homology with PvDBPII (19). Naturally acquired antibodies to both PvDBPII and PvEBPII are associated with protection against *P. vivax* (20–22).

In case of *P. falciparum*, interaction of the parasite ligand PfRH5 with its receptor basigin on the surface of RBCs is essential for invasion (23). PfRH5 is part of a multi-protein complex, referred to as the PCRCR compex, that includes, PfCyRPA, PfRIPR, PfCSS and PfTRAMP. The PCRCR complex is anchored to the merozoite surface by PfTRAMP, which has a transmembrane domain (23–28). Members of the PCRCR complex are localized in apical merozoite organelles referred to as micronemes and rhoptries. While *P. vivax* and the related simian malaria parasite species, *P. knowlesi*, have genes that encode homologs of PfCyRPA, PfRIPR, PfCSS and PfTRAMP, genes encoding PfRH5 homologs are missing in these *Plasmodium* species (29–30). The repertoire of *P. vivax* merozoite proteins that mediate reticulocyte invasion is thus not completely understood. Here, we queried the *Plasmodium* genome database (PlasmoDB) to identify novel *P. vivax* proteins that are expressed in late blood stages, are essential for growth and have the characteristics that suggest the potential to play a role in invasion. We describe the functional analysis of two novel *P. vivax* merozoite proteins, PvMP45 and PvMP36 (PlasmoDB Gene IDs PVX_083375 and PVX_086190), and their *P. knowlesi* orthologs, PkMP45 and PkMP36 (PlasmoDB Gene IDs PKNH_1251300 and PKNH_1353200). We analyzed the potential functional roles of these proteins in host cell invasion and explored their potential as blood stage vaccine candidates for *P. vivax* malaria in combination with previously identified invasion ligands, PvDBP and PvEBP. We also evaluated if antibodies to these merozoite antigens were acquired through natural exposure and whether specific antibody types are associated with protective immunity to *P. vivax*.

## RESULTS

### Expression of *P. vivax* merozoite proteins, PvMP45 and PvMP36, in *P. vivax* blood stages and localization in merozoites

Genome-wide *piggyBac* transposon mutagenesis in *P. knowlesi* has identified PKNH_1251300 (PkMP45) and PKNH_1353200 (PkMP36) as essential genes for survival of *P. knowlesi* blood stage parasites (31). The Modified Mutagenesis Index Score (MMIS), which is a measure of the tolerance of a gene to disruption, has values of 0.178 and 0.401 for PkMP45 and PkMP36 respectively. These low values fall within the threshold for essential genes. A parallel *piggyBac* mutagenesis study in *P. falciparum* demonstrated the essentiality of their *P. falciparum* orthologs, PfMP45, encoded by PF3D7_1404900 (MMIS: 0.738, Mutagenesis Fitness Score (MFS): −3.131), and, PfMP36, encoded by PF3D7_1349600 (MMIS: 0.228, MFS: −3.204) (32). The negative MFS values validate their critical role in parasite survival. Analysis of gene expression based on RNA-Seq data from *P. falciparum* blood stages indicates that the genes encoding PfMP45 and PfMP36 are transcribed in late *P. falciparum* blood stage schizonts and merozoites. The transcription profiles for PfMP45 and PfMP36 were extracted from the *P. falciparum* transcription profile dataset available in PlasmodDB (33) (Fig. S1A). Proteomics data derived from PlamoDB, also confirms that PfMP45 and PfMP36 are expressed in *P. falciparum* blood stages (33). Transcription of genes, PKNH_1251300 and PKNH_1353200, which encode PkMP45 and PkMP36, respectively, was examined by RT-PCR. Both PkMP45 and PkMP36 are transcribed in *P. knowlesi* blood stage schizonts (Fig. S1B). PfMP45 and PfMP36 and their orthologs from *P. vivax* (PVX_083375 and PVX_086190, respectively), *P. knowlesi* (PKNH_1251300 and PKNH_1353200, respectively) and *P. cynomolgi* (PcyM 1206600 and PcyM 1351100, respectively) share high sequence homology with mean pairwise identity of 99.40% for PvMP45 and 99.81% for PvMP36 (Fig. S2). Analysis of PvMP45 and PvMP36 amino acid sequences from 18 diverse *P. vivax* field strains indicates that these proteins are highly conserved (Fig. S3). Bio-informatic analysis reveals that PvMP45 lacks a signal peptide, whereas PvMP36 has a signal peptide that spans amino acids 1 to 19 (Fig. 1A and Fig. S4). Sequence analysis with TMHMM, DeepTMHMM and SMART did not detect presence of a transmembrane domain (TM) or glycophosphatidyl inositol (GPI) anchor (34, 35, 36). Structural analysis using Phyre-2 (37), indicates that PvMP45 and PvMP36 contain multiple ⍺-helices as well as low-complexity regions with disordered loops (Fig. S4).

**FIG 1.**
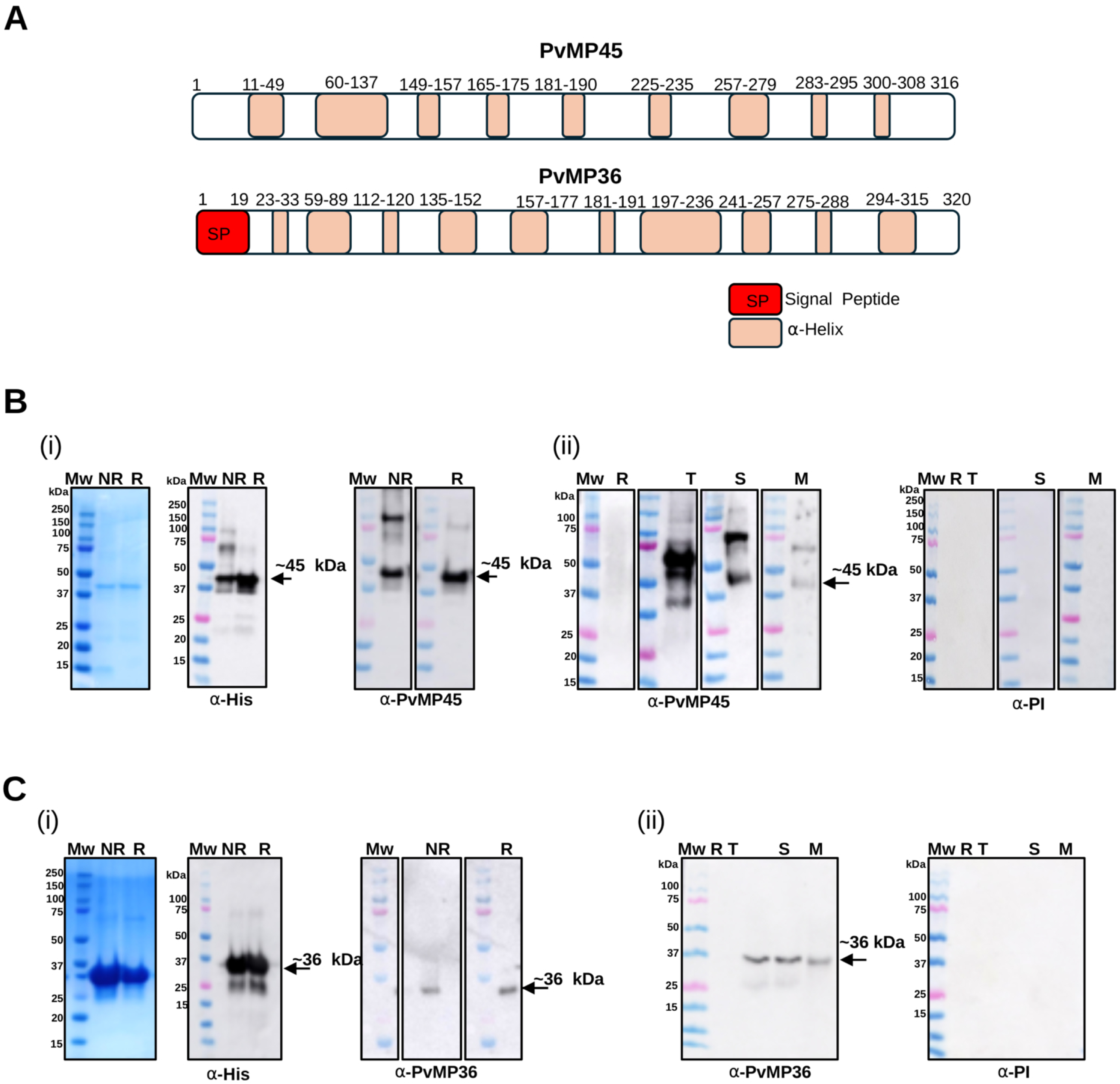
PvMP 45 and PvMP36 expression in blood stages. (A) Schematic representation of primary structure of PvMP45 and PvMP36. Numbers represent amino acid resdues. (B i) Western blot analysis of recombinant PvMP45 under non-reducing (NR) and reducing (R) conditions, probed with anti-His tag antibodies and anti-PvMP45 mouse sera. (B ii) Detection of PkMP45 in *P. knowlesi* blood stages using anti-PvMP45 mouse sera. (C i) Western blot analysis of recombinant PvMP36 under NR and R conditions, probed with anti-His tag antibodies and anti-PvMP36 mouse sera. (C ii) Detection of native PkMP36 in *P. knowlesi* blood stage using anti-PvMP36 mouse sera. R, T, S, and M, *P. knowlesi* Rings, Trophozoites, Schizonts and Merozoites. ⍺-PI, pre-immune mouse sera. Mw, molecular weight markers.

Recombinant PvMP45 and PvMP36 proteins produced in *E. coli* were purified and used to raise antisera in mice and rabbits (Fig. 1B, 1C). Anti-PvMP45 and anti-PvMP36 mouse sera detect expression of the *P. knowlesi* orthologs in lysates of *P. knowlesi* trophozoites, schizonts and merozoites by western blotting due to the high sequence homology (Fig. 1B, 1C). Anti-PvMP45 and anti-PvMP36 mouse sera were also used to localize the proteins in *P. knowlesi* and *P. vivax* blood stages by IFA (Fig. 2 and Fig. 3). Both PkMP45 and PkMP36 have a punctate distribution in *P. knowlesi* schizonts and localize at the apical end of merozoites in mature schizonts (Fig. 3A). PkMP45 and PkMP36 are detected at the apical end of both permeabilized and non-permeabilized *P. knowlesi* merozoites indicating that they translocate to the surface of free merozoites (Fig. 3B, Fig. S5). Both PkMP45 and PkMP36 co-localize with PkDBP in the micronemes of *P. knowlesi* merozoites in mature schizonts (Fig. 3A) as well as free *P. knowlesi* merozoites (Fig. 3B). PvMP45 and PvMP36 also co-localize with PvDBP in late-stage *P. vivax* schizonts confirming their presence in the micronemes (Fig. 3C). Moreover, PvMP45 and PvMP36 co-localize with each other at the apical end of *P. vivax* merozoites confirming that they are localized in the same organelle (Fig. 3C). Use of mouse sera raised against PkRhopH2, which cross-reacts with *P. vivax* rhoptry marker PvRhopH2, confirms that PvMP45 and PvMP36 are not present in the rhoptries (Fig. S6). PvEBP was also detected in *P. vivax* schizonts with antibodies raised against recombinant PvEBPII (Fig. 3D, Fig. S7). Moreover, PvEBP co-localizes with PvDBP, which confirms that PvEBP is also localized in the micronemes (Fig. 3D).

**FIG 2.**
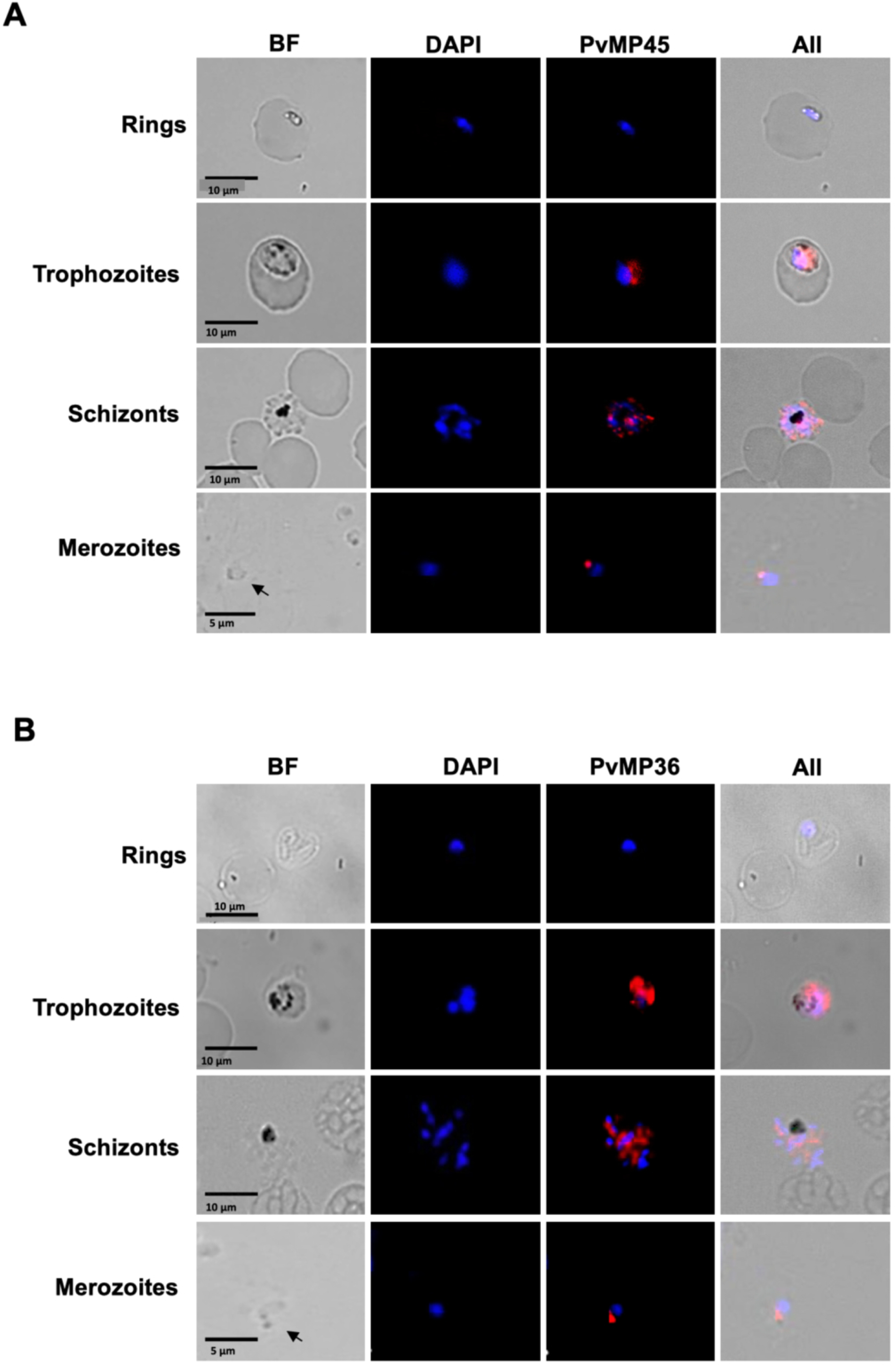
Localization of PkMP45 and PkMP36 in *P. knowlesi* blood stages. (A) Anti-PvMP45 mouse sera detected expression of PkMP45 (red) in trophozoites, schizonts and merozoites. (B) Anti- PvMP36 sera detected expression of PkMP36 (red) in trophozoites, schizonts and merozoites. Bright field (BF) and DAPI (4’,6’-diamidino-2-phenylindole) staining were used to visualize parasite morphology and nuclei, respectively.

**FIG 3.**
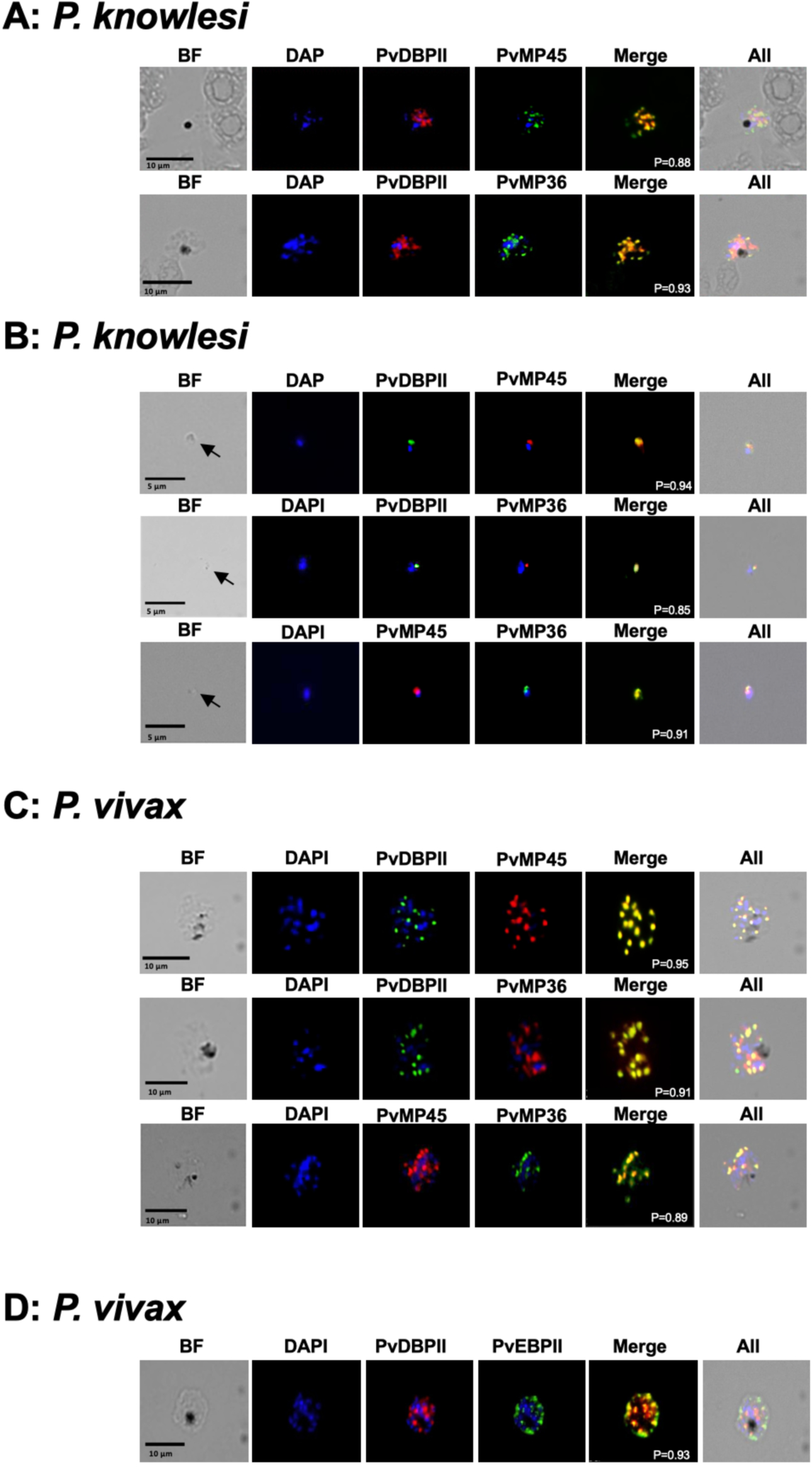
Subcellular localization of PvMP45, PvMP36, and PvEBPII in *P. knowlesi* and *P. vivax* schizonts and merozoites. (A) PkMP45 and PkMP36 were detected in *P. knowlesi* schizonts using anti-PvMP45 and anti-PvMP36 mouse sera (red) respectively. PvDBPII was detected in *P. knowlesi* schizonts using anti-PvDBPII rabbit sera (green). (B) PkMP45 and PkMP36 were detected in *P. knowlesi* merozoites using specific anti-PvMP45 and anti-PvMP36 mouse sera (red) respectively. PvDBPII was detected in *P. knowlesi* merozoites using anti-PvDBPII rabbit sera (green). PvMP45 and PvMP36 were co-localized in merozoites using anti-PvMP45 mouse sera (red) and anti- PvMP36 rabbit sera (green). (C) PvMP45 and PkMP36 were detected in *P. vivax* schizonts using anti-PvMP45 and anti-PvMP36 mouse sera (red) respectively. PvDBPII was detected in *P. vivax* schizonts using anti-PvDBPII rabbit sera (green). (D) Co-localization of PvEBPII and PvDBPII in *P. vivax* schizonts using anti-PvEBPII mouse sera (red) and anti-PvDBPII rabbit sera (green). DAPI, 4’,6’-diamidino-2-phenylindole. P, Pearson’s coefficient values were calculated from average of five images for each panel to quantitate overlap of red and green signals . Pre-immune sera from mice and rabbits did not show any reactivity with schizonts or merozoites.

### Binding of PvMP45, PvMP36 and PvEBPII with human RBCs and reticulocytes

Since several merozoite invasion ligands localized to micronemes, such as PvDBPII, bind host receptors to mediate invasion, we tested whether PvMP45 and PvMP36, as well as the PvDBP homolog, PvEBP, also bind RBCs and/or reticulocytes. Recombinant PRDX6, a cytosolic peroxiredoxin enzyme from human RBCs (38), was used as a negative control and PvDBPII was used as a positive control in the binding assays. PRDX6 did not bind Duffy positive or Duffy negative RBCs or reticulocytes (Fig. 4). As expected, PvDBPII bound Duffy positive RBCs and reticulocytes but did not bind Duffy negative RBCs or reticulocytes (Fig. 4). PvEBPII, PvMP45 and PvMP36 bound both Duffy positive and Duffy negative RBCs as well as reticulocytes (Fig. 4) indicating that they do not bind the Duffy antigen but bind alternative receptors that are present on RBCs and reticulocytes.

**FIG 4.**
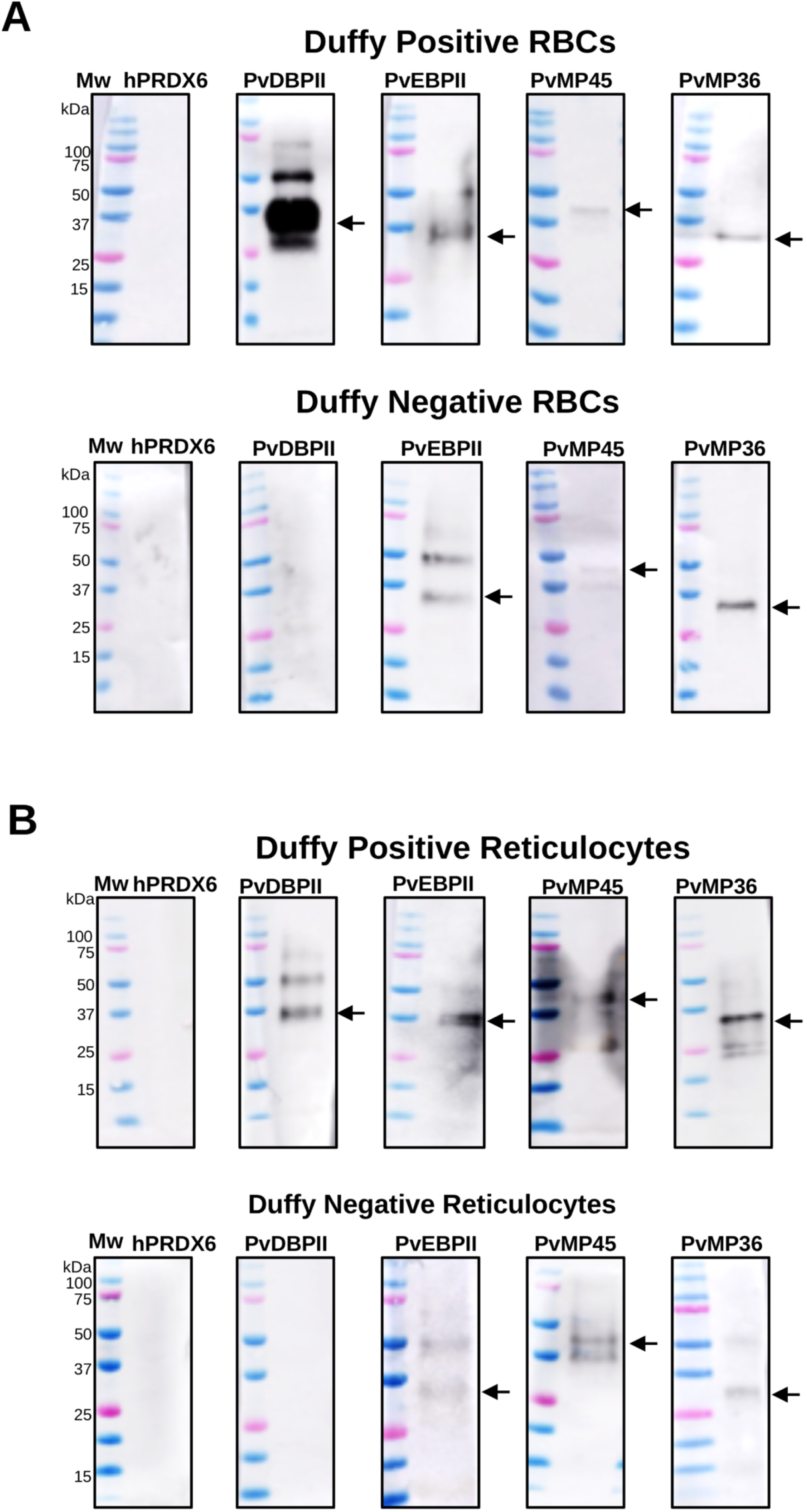
Binding of PvMP45, PvMP36, PvEBPII and PvDBPII with RBCs and reticulocytes. (A) Binding of PvMP45, PvMP36, PvEBPII and PvDBPI with Duffy-positive and Duffy-negative human RBCs. (B) Binding of PvMP45, PvMP36, PvEBPII and PvDBPI with Duffy-positive and Duffy-negative human reticulocytes. Human PRDX6 was used as a negative control. Arrows indicate expected size of bound proteins.

### Ability of antibodies raised against PvMP45, PvMP36, PvEBPII and PvDBPII to inhibit *P. vivax* invasion of reticulocytes

Given the potential role of PvMP45, PvMP36, PvEBPII and PvDBPII to bind host receptors during merozoite invasion of reticulocytes, we examined if antibodies raised against these proteins can inhibit *P. vivax* invasion *in vitro* (Fig. 5). IgGs purified from pre-immune rabbit sera did not show any significant inhibition (<15%) compared to controls without any IgGs (Fig. 5). Rabbit IgGs raised against PvDBPII were the most potent invasion inhibitory antibodies with ∼80% inhibition at IgG concentration of 5 mg/ml. Rabbit IgGs raised against PvEBPII, PvMP45 and PvMP36 also had invasion inhibitory activity (∼50% inhibition at IgG concentration of 5 mg/ml) but were less potent compared to anti-PvDBPII rabbit IgGs (Fig. 5A). Combining IgGs raised against PvDBPII with IgGs against PvMP36 had a statistically significant additive invasion inhibitory effect (Fig. 5B).

**FIG 5.**
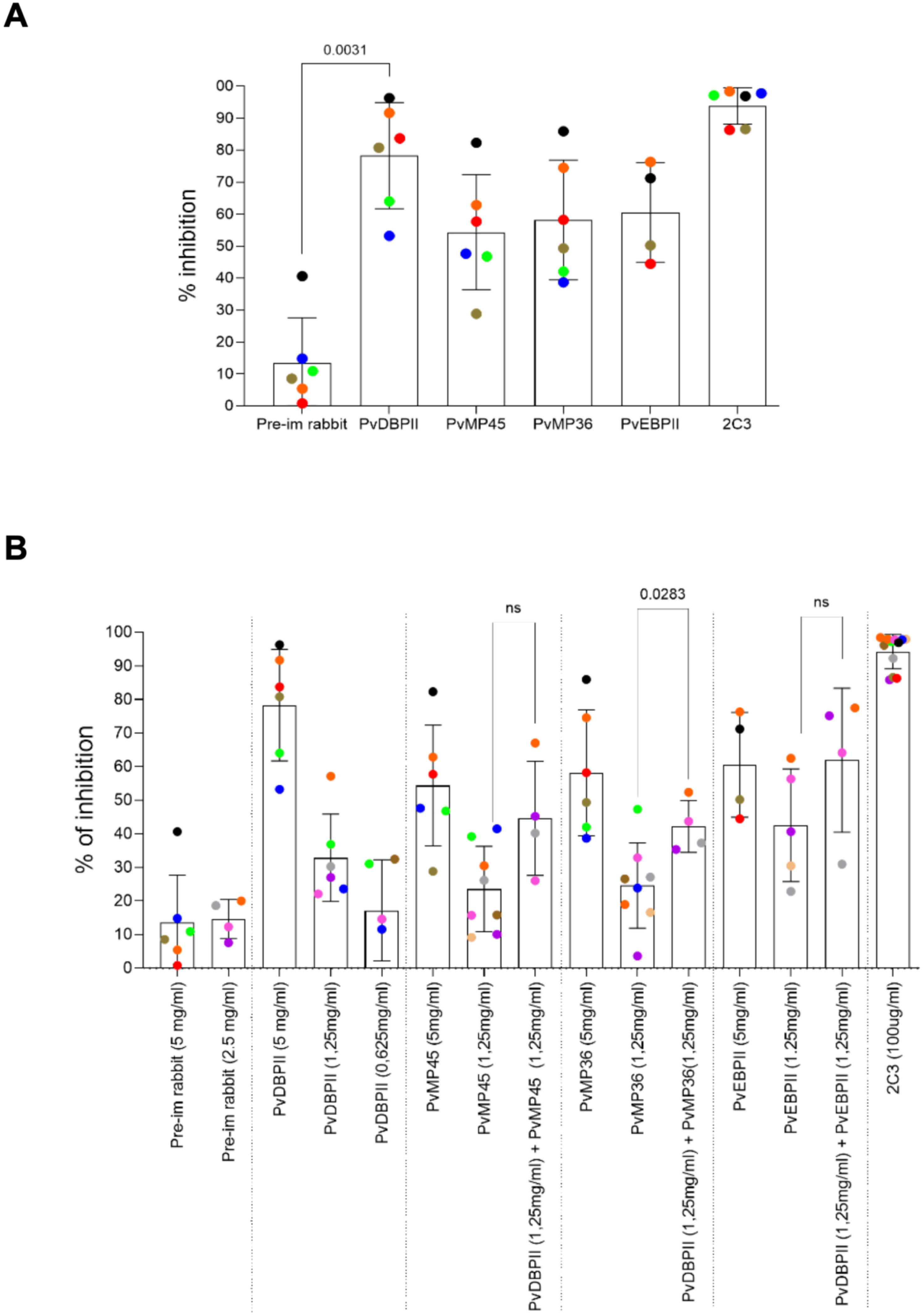
Inhibition of reticulocyte invasion by *P. vivax* with antibodies to PvMP45, PvMP36, PvEBPII and PvDBPII. (A) Purified IgGs from rabbit sera raised against PvMP45, PvMP36, PvEBPII and PvDBPII were tested for inhibition of reticulocytes by *P. vivax*. Anti-PvDBPII showed the strongest inhibition (∼80% at 5 mg/ml), while antibodies to PvMP45, PvMP36, and PvEBPII showed moderate inhibition (∼50%). Pre-immune IgGs had minimal inhibitory effect (∼10%). Inhibitory mouse monoclonal antibody 2C3 against DARC was included as a positive control. (B) Rabbit IgGs against PvDBPII, PvMP45, PvMP36 and PvEBPII were tested for inhibition of reticulocyte invasion individually at different concentrations (5mg/ml, 2.5mg/ml and 1.25mg/ml) and in combination. PvDBPII IgG showed potent inhibition in a dose-dependent manner, while combination assays of PvDBPII with PvMP45, PvMP36, or PvEBPII revealed additive effects. However, additive effect of only PvDBPII + PvMP36 combination reached statistical significance (P value = 0.0283). ns, non-significant P-values.

### PvMP45 and PvMP36 interact with other invasion related proteins on the merozoite surface

Several parasite proteins that play a role in invasion form protein complexes on the merozoite surface (26–30, 39, 40). We examined if PkMP45 and PkMP36 also interact with other invasion-related proteins to form protein complexes on the surface of *P. knowlesi* merozoites. Following isolation of *P. knowlesi* merozoites, the surface proteins were crosslinked and the merozoite lysate was used for immunoprecipitation with rabbit antibodies to PvMP45 and PvMP36. Western blot analysis of the immunoprecipitates confirmed successful pull-down of PkMP45 and PkMP36 (Fig. S8). Proteins that are pulled down by immunoprecipitation with anti-PvMP45 and anti-PvMP36 rabbit sera were identified by LC-MS/MS (Supplementary Tables S1 and S2). Proteins found in immunoprecipitates with anti-PvMP45 and anti-PvMP36 sera that meet the following criteria, (i) consistent detection across five biological replicates; (ii) a fold change >1.3 in PvMP45 or PvMP36 pull-downs compared to control pull downs with pre-immune sera with an adjusted P-value of <0.05; and (iii) known or predicted roles in erythrocyte invasion or known localisation on the merozoite surface or apical organelles like micronemes and rhoptries, are shown in Tables 1 and 2. In immunoprecipitates with anti-PvMP45 sera, this analysis identifies peptides corresponding to PkMP45 as well as the following invasion related proteins: PkMSP1, PkRAP1, PkRAP2, PkEMP, PkMSP7-like protein, and PkMSP-7D (Table 1). These peptides were significantly enriched in anti-PkMP45 immunoprecipitation eluates as compared to pre-immune rabbit sera. In case of immunoprecipitation with anti-PkMP36 rabbit sera, LC-MS/MS analysis detected presence of PkMP36 as well as PkMSP1, PkRAP1, PkRhopH3, PkMSA180, PkRAP2, PkEMP, PkMSP7D-like protein and putative rhoptry protein PKNH_0316800 (Table 2). Several proteins in the PkMP45 and PkMP36 immunoprecipitates were common, but others were unique.

**TABLE 1.**
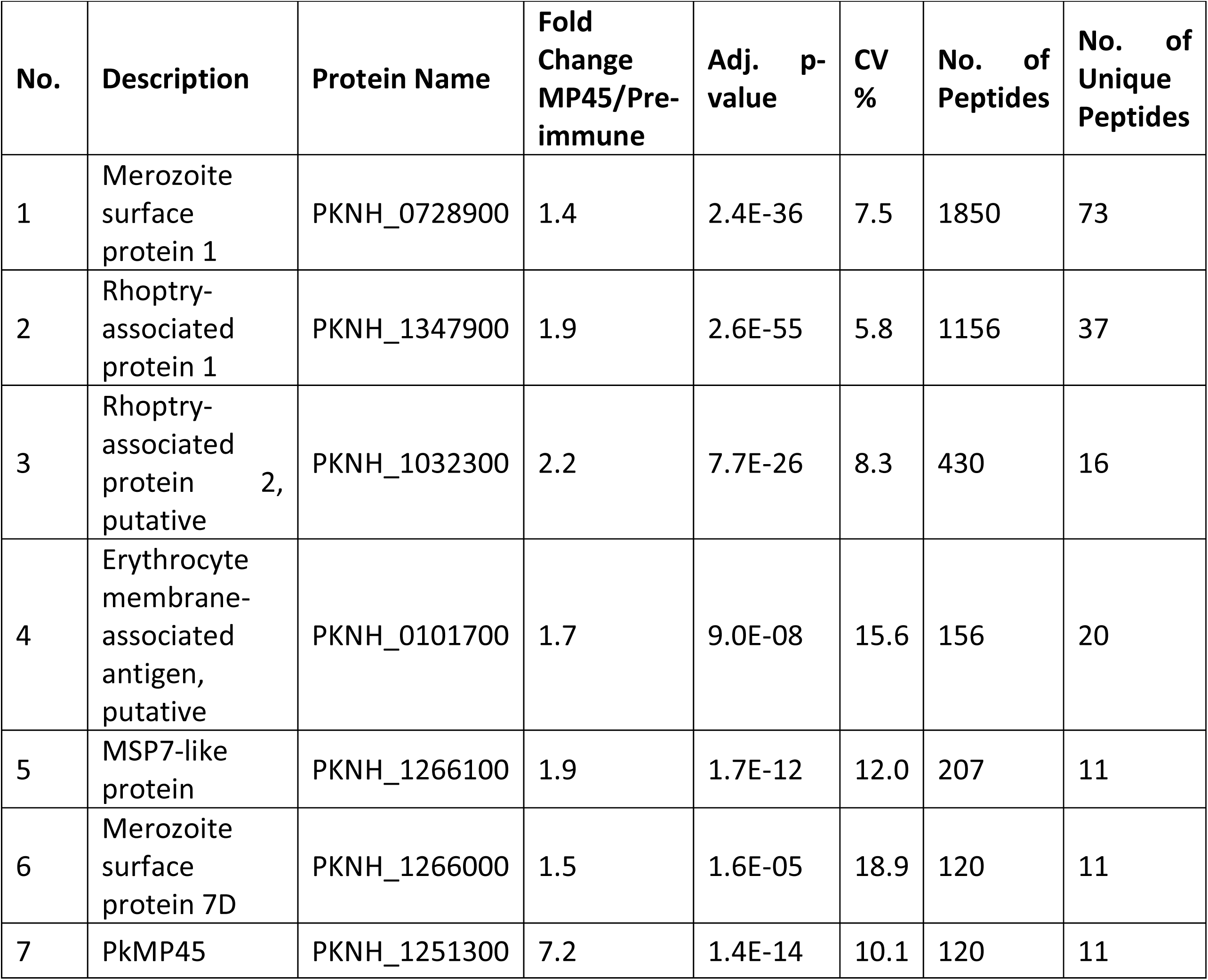
PkMP45 associated proteins in *P. knowlesi* merozoites.

| No. | Description | Protein Name | Fold Change MP45/Pre-immune | Adj. p-value | CV % | No. of Peptides | No. of Unique Peptides |
| --- | --- | --- | --- | --- | --- | --- | --- |
| 1 | Merozoite surface protein 1 | PKNH_0728900 | 1.4 | 2.4E-36 | 7.5 | 1850 | 73 |
| 2 | Rhoptry-associated protein 1 | PKNH_1347900 | 1.9 | 2.6E-55 | 5.8 | 1156 | 37 |
| 3 | Rhoptry-associated protein 2, putative | PKNH_1032300 | 2.2 | 7.7E-26 | 8.3 | 430 | 16 |
| 4 | Erythrocyte membrane-associated antigen, putative | PKNH_0101700 | 1.7 | 9.0E-08 | 15.6 | 156 | 20 |
| 5 | MSP7-like protein | PKNH_1266100 | 1.9 | 1.7E-12 | 12.0 | 207 | 11 |
| 6 | Merozoite surface protein 7D | PKNH_1266000 | 1.5 | 1.6E-05 | 18.9 | 120 | 11 |
| 7 | PkMP45 | PKNH_1251300 | 7.2 | 1.4E-14 | 10.1 | 120 | 11 |

**TABLE 2.** PkMP36 associated proteins in *P. knowlesi* merozoites.

| No. | Description | Protein Name | Fold Change MP36/Pre-immune | Adj. p-value | CV % | No. of Peptides | No. of Unique Peptides |
| --- | --- | --- | --- | --- | --- | --- | --- |
| 1 | Merozoite surface protein 1 | PKNH_0728900 | 1.3 | 1.20E-26 | 8.8 | 1944 | 78 |
| 2 | Rhoptry-associated protein 1 | PKNH_1347900 | 1.6 | 3.71E-30 | 8.1 | 1210 | 44 |
| 3 | High molecular weight rhoptry protein 3, putative | PKNH_0703100 | 1.4 | 8.04E-13 | 12.4 | 512 | 26 |
| 4 | Merozoite surface protein MSA180, putative | PKNH_0814000 | 1.3 | 0.000139 | 21.5 | 344 | 32 |
| 5 | Rhoptry-associated protein 2, putative | PKNH_1032300 | 1.5 | 2.25E-06 | 17.9 | 462 | 18 |
| 6 | PkMP36 | PKNH_1353200 | 48.0 | 1.21E-135 | 2.2 | 359 | 19 |
| 7 | Erythrocyte membrane-associated antigen, putative | PKNH_0101700 | 1.6 | 3.68E-05 | 19.6 | 166 | 21 |
| 8 | Merozoite surface protein 7D | PKNH_1266000 | 1.4 | 3.92E-04 | 22.1 | 205 | 11 |
| 9 | Rhoptry protein, putative | PKNH_0316800 | 1.4 | 0.03 | 33.7 | 121 | 8 |

To validate the association of the rhoptry protein PkRAP1 with PkMP45 and PkMP36, we performed immunoprecipitation of *P. knowlesi* merozoite lysates with anti-PvMP45, anti- PvMP36, and anti-PkRAP1 sera and verified the presence of interacting partners by western blotting. Western blot analysis revealed that immunoprecipitation with anti-PvMP45 rabbit sera pulls down PkMP36 and PkRAP1 along with PkMP45 (Fig. 6A). Similarly immunoprecipitation using anti-PvMP36 antibodies successfully pulls down not only PkMP36 but also PkMP45 and PkRAP1 (Fig. 6A). In case of immunoprecipitation with anti-PkRAP1 antibodies, PkRAP-1 as well as PkMP36 were detected in the immunoprecipitates but PkMP45 was not detected (Fig. 6A). The amount of PkMP45 in the immunoprecipitate with anti-PkRAP1 antibodies may be below the limit of detection due to weaker binding compared to binding to PkMP36. The binding of PkRAP1 to PvMP45 and PvMP36 was also directly tested in an ELISA-based protein-protein interaction assay. In this assay, 96-well ELISA plates were coated with recombinant PvMP45 and PvMP36 followed by the addition of increasing concentrations of PkRAP1. Binding of PkRAP1 to the different coated receptors was analyzed using anti-PkRAP1 sera. Recombinant PvDBPII was used as negative control. Concentration-dependent binding was observed between the coated proteins PvMP45 and PvMP36 and PkRAP1 (Fig. 6B). PvMP36 displayed stronger binding to PkRAP1 compared to PvMP45.

**FIG 6.**
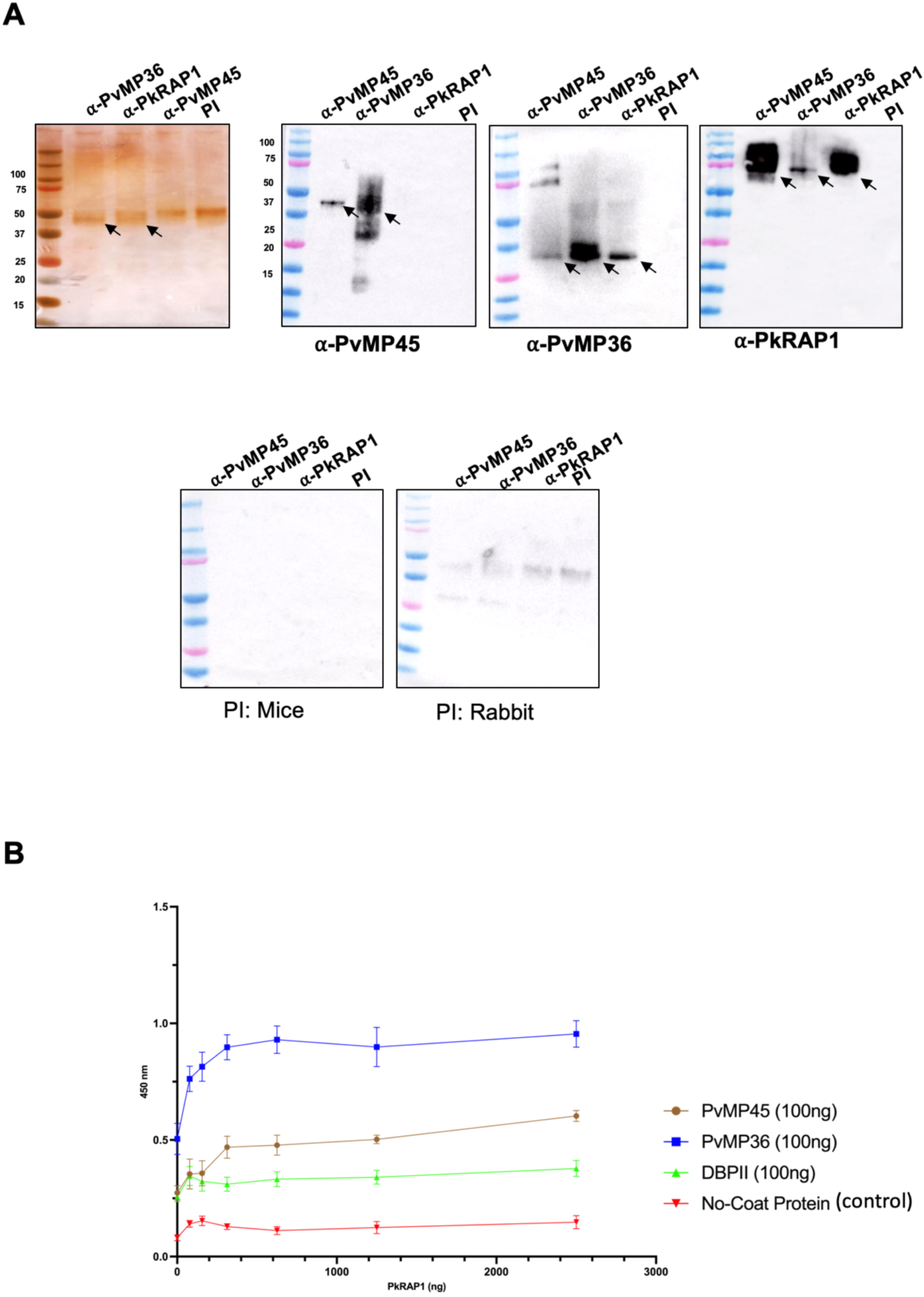
Co-Immunoprecipitation and ELISA-Based protein interaction assays. (A) *P. knowlesi* merozoites were crosslinked, lysed and used for immunoprecipitation with anti-PvMP45, anti- PvMP36, anti-PkRAP1 mouse sera and pre-immune (PI) mouse sera as control. Immunoprecipitates were separated by SDS-PAGE gel electrophoresis and detected by silver staining. Immunoprecipitation of specific proteins was detected by western blotting using antisera against PkMP45, PkMP36 and PkRAP1. (B) ELISA plate-based analysis of binding of PkRAP1 with PkMP45 and PkMP36. A concentration dependent specific binding interaction was observed between PkRAP1 and PkMP36 as well as PkRAP1 and PkMP45. PkRAP1 showed higher binding with PkMP36 than PkMP45. PvDBPII showed no interaction with PkRAP1. Bars represent the SEM from three independent experiments.

### Naturally acquired antibodies against PvMP45, PvMP36, PvEBPII and PvDBPII and association with protection against *P. vivax* malaria

Sera from children residing in a *P. vivax* endemic region of Papua New Guinea (PNG) were analyzed for presence of naturally acquired antibodies against the novel microneme proteins PvMP45 and PvMP36 as well as PvDBPII and PvEBPII (Fig. S9). Significant IgG, IgG1 and IgG3 responses were evident against both PvMP45 and PvMP36 in PNG children compared to controls, particularly against PvMP36. Antibodies against PvMP45 and PvMP36 bound FcyRI and FcyRIIa, which are found on monocytes/activated neutrophils, and FcyRIIIa, which is found on monocytes, macrophages, NK cells and neutrophils (Fig. S10). Both PvMP45 and PvMP36 antibodies induced significant fixation of C1q (Fig. S10). Total IgG antibodies against PvMP36 but not PvMP45 were associated with protection against clinical vivax malaria (Fig. 7A, 7B). IgGs to PvEBPII had the strongest association with protection followed by PvMP36 and PvDBPII. IgGs to PvM45 and PvCSP did not have any significant association with protection. Similar patterns were observed for IgG1, although IgG1 to PvDBPII Sal1 was not significantly associated with protection. All antigens induced IgG3 antibodies that were associated with protection from clinical P*. vivax* episodes, with PvEBPII and PvMP36 having the strongest associations. No antigens elicited C1q-fixing antibodies that were associated with protection (Fig. 7C, 7D). Since anti-PvEBPII FcyR-binding antibodies were consistently associated with protection, phagocytic activity against opsonized PvEBPII-coated beads was evaluated. Plasma from *P. vivax*- positive samples displayed strong opsonizing activity with PvEBPII-coated beads compared to very low opsonizing activity when beads were incubated with control plasma (Fig. 7E). Phagocytic activity positively correlated with presence of FcyRI-binding antibodies in the same samples (r=0.787) (Fig. 7F).

**FIG 7.**
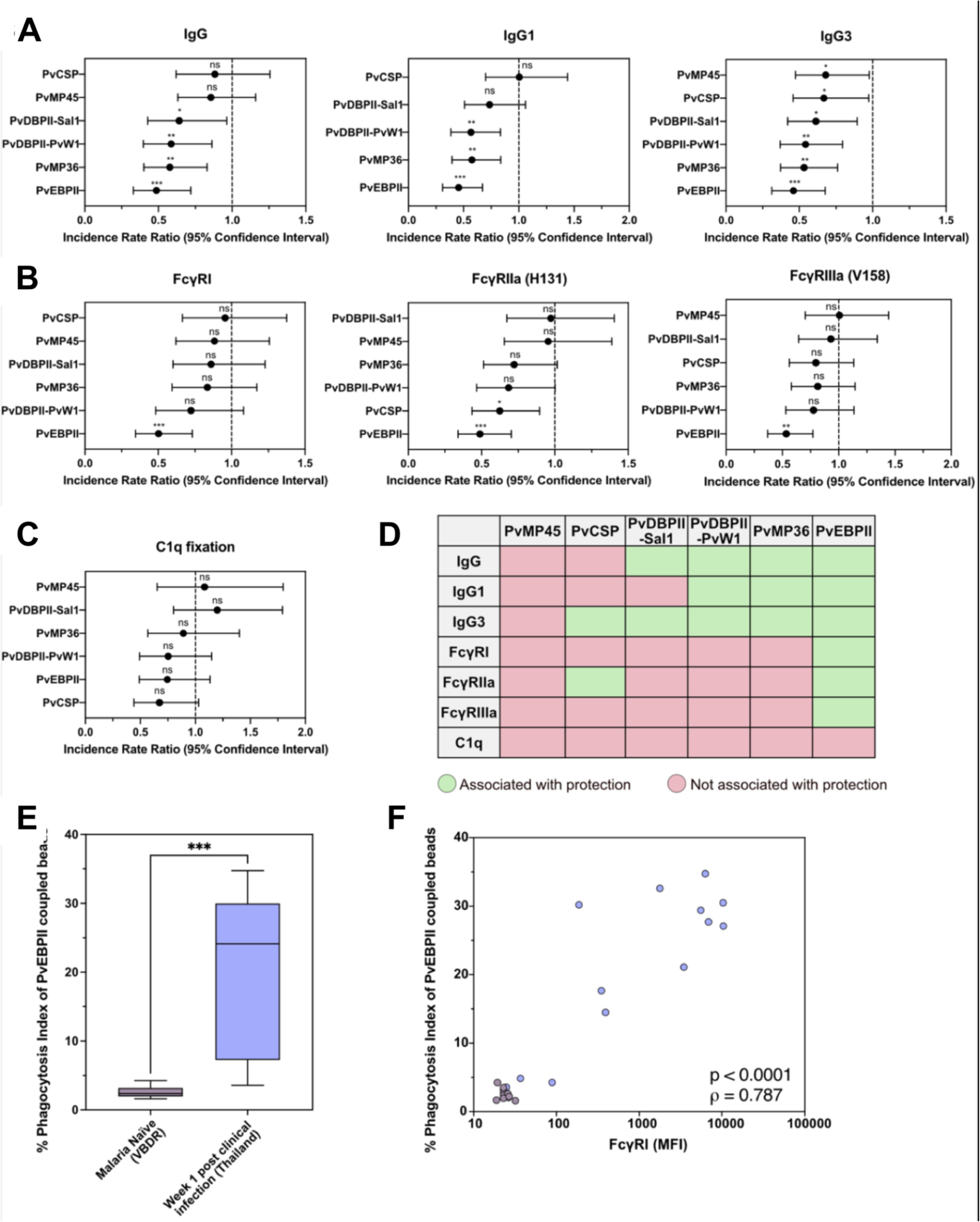
Association between antibody responses to *P. vivax* proteins and protection against *P. vivax* malaria in Papua New Guinean children. (A) Association between IgG, IgG1 and IgG3 antibody levels to PvCSP, PvDBPII-PvW1 and PvDBPII-SalI, PvEBPII, PvMP36 and PvMP45 and protection against clinical *P. vivax* malaria in Papua New Guinean children aged 1 – 3 years old (n = 187) residing in a *P. vivax* endemic region as represented by incidence rate ratios (IRRs), 95% confidence intervals and P values derived from a generalized estimation equation (GEE) model comparing high versus low tertiles of responders. PvMP45 was excluded from IgG1 analysis as tertile groups could not be distinguished. (B) Assocation between FcγRI, FcγRIIa (H131) and FcγRIIIa (V158) dimer binding to *P. vivax* antigen-specific antibodies and protection against *P. vivax* malaria, represented by IRRs. (C) Association between C1q fixation levels and protection against *P. vivax* malaria, represented by IRRs. (D) Summary of protective associations between IgG levels and functional antibody levels and risk of *P. vivax* malaria incidence (green = associated with protection; pink = not associated with protection). (E) Phagocytisis of PvEBPII coupled beads opsonized with human sera from malaria naïve cohort samples from Melbourne, Victoria, Australia (Victorian Biospecimen Donor Registry (VBDR)) (n = 11) and samples one week post clinical *P. vivax* infection from Tha Song Yang, Thailand (n = 12) measured by flow cytometry. (F) Correlation of phagocytosis activity with FcyRIa binding via Luminex (Mean fluorescence intensity) calculated using Spearman’s rank correlation tests. For statistical significance p-values were represented as <0.0001 (****), ≤ 0.0002 (***), ≤ 0.0021 (**),≤0.0332 (*), > 0.05 (ns).

## DISCUSSION

Although significant progress has been made in understanding RBC invasion by *P. falciparum*, the invasion process of *P. vivax* remains less well characterized due to lack of a continuous *in vitro* culture system. To date, only few *P. vivax* ligands that mediate host cell entry have been identified. In the present study, we have identified and characterized two novel *P. vivax* proteins, PvMP45 and PvMP36, which are expressed in blood stages including merozoites. Their essentiality is strongly supported by genome-wide piggyBac transposon mutagenesis studies in *P. knowlesi* and *P. falciparum*, showing that their orthologs in both species are refractory to disruption (31, 32). This functional indispensability, along with their conservation across 18 geographically diverse *P. vivax* field isolates (Fig. S3), highlights their potential as novel vaccine targets. PvMP45 and PvMP36 and their *P. knowlesi* orthologs, PkMP45 and PkMP36, are localized in the micronemes (Fig. 3) suggesting that they may play a role in invasion. Indeed, PvMP45 and PvMP36 bind RBCs and reticulocytes (Fig. 4) suggesting that they may mediate receptor binding during invasion.

Parasite invasion ligands are often present in protein complexes on the merozoite surface (26–30, 39, 40). Immunoprecipitation experiments indicate that both PvMP45 and PvMP36 may form invasion protein complexes on the merozoite surface. Immunoprecipitation with anti-PvMP45 sera identified the following invasion related interacting partners: PkMSP1, PkRAP1, PkRAP2, PkMSP7-like proteins and PkMSP7D, which are implicated in host cell invasion (Table 1). Similarly, immunoprecipitation with anti-PvMP36 sera identified the following invasion related interacting partners: PkMSP1, PkRAP1, PkRAP2, PkRhopH3, PkMSP180, PkMSP7D and a putative rhoptry protein PKNH_0316800 (Table 2). Among the interacting proteins identified, only recombinant PkRAP1 and anti-PkRAP1 sera were available for use in validation experiments. Co-immunoprecipitation experiments with antisera to PvMP45, PvMP36 and PkRAP1 confirmed interaction of PvMP36 and PvMP45 with PkRAP1 (Fig. 6). In addition, direct binding experiments using recombinant PkMP36, PkMP45 and PkRAP1 confirmed that PkMP36 and PkMP45 bind PkRAP1.

The essentiality of PkMP45 and PkMP36 and their localization in the micronemes with PvDBP and PvEBP suggests that PvMP45 and PvMP36 play an important role in reticulocyte invasion. However, though PvMP45 and PvMP36 bind Duffy negative RBCs and reticulocytes, these interactions are not sufficient to enable *P. vivax* merozoite infection of Duffy negative RBCs or reticulocytes. These parasite ligands may only support reticulocyte invasion by the Duffy pathway mediated by PvDBP. Given the central role of PvDBP in reticulocyte invasion by *P. vivax* merozoites, it is not surprising that anti-PvDBPII IgGs are most efficient at blocking *P. vivax* invasion compared to IgGs targeting PvMP45, PvMP36 and PvEBPII. The combination of IgGs against PvMP36 and PvEBPII yields an additive inhibitory effect in *P. vivax* invasion assays (Fig. 5).

Sero-epidemiological analysis conducted in a longitudinal cohort of 1-4 year old children residing in a *P. vivax* endemic region of PNG found that naturally acquired IgGs to PvEBPII, PvDBPII and PvMP36 were associated with protection from *P. vivax* malaria (Fig. 7). The strongest associations with protection were observed for IgG, IgG1, and IgG3 to PvEBPII, PvDBPII, and PvMP36. Antibodies to PvMP45 were not associated with protection against *P. vivax* malaria (Fig. 7). Antibodies to PvEBPII displayed the highest Fcγ receptor (FcγRI, FcγRIIa and FcγRIIIa) binding (Fig. 7). Importantly, Fcγ receptor binding by antibodies against PvEBPII was associated with protection suggesting that cytophilic antibody responses may play an important role in protection via antibody dependent cell mediated responses (Fig. 7). Indeed, a beed-based assay confirmed that naturally acquired human antibodies against PvEBPII could mediate opsonic phagocytosis in an *in vitro* assay using human monocytes (Fig. 7). Our understanding of the role of Fcy-receptor mediated functions against merozoites in immunity to *P. vivax*, and malaria more generally, is limited. Studies have suggested that opsonic phagocytosis by monocytes and neutrophils, and ADCC by NK cells and neutrophils may be important mechanisms (41, 42, 43). Fc-y receptor binding by antibodies has been shown to protect against blood stages in mouse models (44).

Observations from the *in vitro* invasion inhibition assays and field studies on immune correlates, suggest that PvDBPII, PvEBPII and PvMP36 may provide an attractive combination of merozoite antigens for development of a multivalent blood stage vaccine for *P. vivax* malaria. Naturally acquired IgG levels against these antigens are clearly associated with protection against *P. vivax* malaria. In addition, antibodies to PvDBPII are potent at blocking *P. vivax* invasion. Combination of antibodies to PvDBPII and PvMP36 could further enhance this invasion inhibitory effect. In addition, antibodies to PvEBPII could invoke an alternative immune mechanism to clear merozoites by antibody dependent cellular mechanisms such as opsonic phagocytosis. The combination of PvDBPII, PvEBPII and PvMP36 could provide a potent combination of *P. vivax* blood stage antigens for development of a multivalent, high efficacy vaccine for *P. vivax* malaria. Next steps in this direction will require better understanding of the diversity of these antigens, further validation of the protective role of these antigens in other cohorts and evaluation of antibodies raised against these antigens for inhibition of a large panel of diverse isolates. Finally, there is a need to identify a delivery platform to deliver a multivalent vaccine candidate to elicit high levels of long lasting efficacy. Such a vaccine could be a key component in a strategy to eliminate *P. vivax* malaria.

## MATERIALS AND METHODS

### *In-vitro P. knowlesi* parasite culture and isolation of viable merozoites

*P. knowlesi* H strain blood stage parasites adapted to grow in human type O, Rh+ Duffy positive blood (kindly provided by Dr. Robert Moon, London School of Hygiene and Tropical Medicine) were cultured as described earlier (45). The protocol for isolating *P. knowlesi* merozoites was adapted from the work of Lyth et al., 2018 (46). Compound 2, a protein kinase was used to inhibit schizont rupture and enable isolation of fully mature segmented schizonts that were mechanically disrupted by repeated pipetting and passed through a 5 μm filter. Cell debris were separated by centrifugation at 1200 g and merozoites were collected from the supernatant by centrifugation at 2400 g.

### Reverse Transcription PCR Analysis

Total RNA was extracted from *P. knowlesi* schizont stage cultures using Trizol reagent (Thermo Fisher Scientific) following the manufacturer’s protocol. Each RNA sample (2 µg) was used to synthesize cDNA using the SuperScript III First-Strand Synthesis System (Thermo Fisher Scientific) as described in Supplementary Text S1.

### Human study population and ethics approval

Plasma samples collected from archival study cohorts from PNG (47) and Thailand (48), along with negative controls from Melbourne (Volunteer Biospecimen Donor Registry, VBDR) were used. Written informed consent or assent was obtained from all participants, and/or their guardians, prior to enrolment. Studies were approved by the Medical Research and Advisory Committee of the Ministry of Health in PNG (MRAC 05.19), the Ethics Committee of the Faculty of Tropical Medicine, Mahidol University, Thailand (MUTM 2014-025-01 and 02), and the WEHI Human Research Ethics Committee (07/07 and 14/02). In PNG, plasma samples were utilized from 187 children collected at the recruitment timepoint, to enable association of measured antibodies with subsequent incidence of clinical vivax infections as described in Supplementary Text S2. In the Thai study, plasma samples were collected from 12 individuals at the week 1 post clinical infection timepoint and used to measure matched antibodies and phagocytic activity as described in Supplementary Text S2.

### Multiplexed Luminex assay for measuring antigen-specific antibodies and capacity for FcyR- binding and complement fixation

To enable measurement of antibody repsonses to multiple *P. vivax* proteins, we utilized a multiplex, magnetic bead-based Luminex assay as previously described (49). Antigen-specific total IgG, IgG1 and IgG3 subclass antibodies were measured by incubation of coupled-beads with plasma (1:100) dilution followed by relevant secondary antibodies as previously described (50). Capacity of IgGs against different antigens for FcyRI, FcγRIIa and FcγRIIIa binding and complement fixation was evaluated as previously described (51).

### Opsonic phagocytosis assay

Ability of PvEBPII to elicit opsonic phagocytosis was evaluated using PvEBPII coupled to amine fluorescent (PE) 2.0 µm beads (Sigma) and undifferentiated THP-1 pro- monocyte cell line (ATCC, USA) as previously described (42, 51). Briefly, diluted plasma samples were incubated with 20 µL of PvEBPII-coupled beads, washed and incubated with THP-1 cells for 40 minutes at 37 °C. The cells were fixed with 2% (v/v) paraformaldehyde and analyzed by FACS on a LSR Fortessa X20b (BD Biosciences). The phagocytosis index (%) was calculated using the percentage of single THP-1 cells that were positive for PE-staining indicating phagocytosis of PvEBPII-coupled fluorescent beads.

### Analysis of association of antibodies with protection from clinical malaria

To test the association between antibodies (total IgG, subclass, functional) with protection from clinical *P. vivax* malaria, we utilized a negative binomial generalized estimating equation (GEE) model as previously described (22). Antibody levels were first categorized into tertiles to separate individuals into high, medium and low groups, with the low group used as the reference in the analysis. Results were presented as incidence rate ratios (IRR), and adjusted for age, season, village of residence and exposure utilizing the molecular force of blood-stage infection.

### Statistical analysis for serology

Statistical analyses were performed using Stata v17.1 (StataCorp, Texas, USA) or R Statistical Sotware. Graphs were generated using either the ggplot2 package in R or GraphPad Prism v10. Wilcoxon rank-sum tests were used to compare IgG and functional antibody levels and Spearman’s rank correlation coefficient was used for assessment of correlations between phagocytosis activity and FcyRI binding.

### Expression of recombinant PvMP45, PvMP36, PvEBPII and PvDBPII proteins

Synthetic genes encoding amino acid (aa) sequences of PvEBPII (aa 171-491), PvMP45 (aa 1-316) and PvMP36 (aa 20 – 321) from *P. vivax* Sal-I strain and optimized for recombinant protein expression in *E. coli* were cloned in the *E. coli* expression vector pET28a (+) (Novagen, USA). *E. coli* BL21 (DE3) strain cells (New England Biolabs, USA) with expression plasmids for either PvMP45 or PvMP36 were cultured to mid-log phase with OD_600_ of 0.6–0.8 and induced with 0.5 mM IPTG at 16 °C overnight to express recombinant PvMP45 and PvMP36. Following induction, the cells were harvested by centrifugation and lysed by sonication. Recombinant PvMP45 and PvMP36 proteins were purified from soluble cytosolic fractions under native conditions by metal affinity chromatography (Ni-NTA) followed by gel permeation chromatography (GPC) using a Superdex 600 column (GE Healthcare, Sweden). Recombinant PvEBPII was expressed in *E. coli* Shuffle 30 cells transformed with pET28a-PvEBPII construct. Recombinant protein expression was induced with 1 mM IPTG at 30℃ for 6 hours and purified from cytosolic fraction by Ni-NTA chromatography followed by GPC using a Superdex 600 column (GE Healthcare, Sweden). Expression of PvDBPII (region II of *P. vivax* Duffy binding protein) has been described earlier (52, 53).

### Raising antibodies against PvMP45, PvMP36 and PvEBPII in mice and rabbits

Recombinant PvMP45, PvMP36, PvDBPII and PvEBPII proteins formulated with Freund’s adjuvant were used to raise antisera in mice and rabbits as described in Supplementary Text S3.

### Western blot analysis of *P. knowlesi* blood stages

Specific antisera raised in mice were used to detect PvMP45 and PvMP36 in lysates of *P. knowlesi* rings, trophozoites, schizonts, and merozoites by western blotting. Pellets of synchronized *P. knowlesi* cultures were harvested at different stages by centrifugation and lysed using RIPA buffer (50 mM Tris, 150 mM NaCl, 1 mM EDTA, 1% NP-40, 5% glycerol (pH 7.4) on ice for 1 hour. Supernatants were collected by centrifugation at 19,500g for 40 minutes at 4°C, separated by SDS-PAGE and used for western blotting using anti-PvMP45, anti-PvMP36, and pre-immune mouse sera using standard techniques.

### Immunofluorescence assay (IFA) to detect expression and localize PvMP45 and PvMP36 in *P. vivax* and *P. knowlesi* blood stage parasites

Immunofluorescence assays (IFAs) were performed with *P. knowlesi* asexual blood stages and *P. vivax* schizonts, which were smeared on glass slides, air dried, and fixed for 30 minutes with pre-chilled acetone:methanol (90:10) as described previously (15). For co-localization studies in *P. knowlesi* asexual stages, anti-PvMP45 and anti- PvMP36 mouse sera (1:50 dilution) were used along with rabbit antisera against microneme proteins, PvEBP (1:200 dilution) and PvDBP (1:300 dilution), or rhoptry protein, PkRhopH2 (1:100 dilution). The slides were incubated with primary antisera for 1h at RT, washed with PBST, and incubated with either Alexa Fluor 488 or Alexa Fluor 594-conjugated secondary anti-mouse IgG or anti-rabbit IgG goat sera for 1 hour. The slides were washed with PBST, mounted in ProLong Gold antifade reagent with 4’,6-diamidino-2-phenylindole (DAPI) (Invitrogen), viewed with a GE DeltaVision Elite fluorescence microscope and the images were analysed using *softWorx*, software.

### Erythrocyte binding assay with recombinant PvMP45, PvMP36, PvEBPII and PvDBPII

Recombinant PvMP45, PvMP36, PvEBPII, and PvDBPII were used in RBC binding assays. Briefly, 10µg of each recombinant protein was incubated with 10µl of packed RBCs for 1 hour at room temperature. The RBCs with bound proteins were separated by centrifugation through a silicone oil (85 % silicone −15 % Nujol) at 3000g for 1 min. Proteins bound to erythrocytes were eluted by incubation with 20 µl of 300mM NaCl at room temperature for 5 minutes, separated by centrifugation and detected by western blotting. Recombinant PvDBPII was used as a positive control and recombinant PRDX6 (human peroxiredoxin 6), a cytosolic RBC enzyme (34), was used as a negative control. The binding assay was performed using RBCs as well as reticulocytes isolated from both Duffy-positive and Duffy-negative blood. A commercially available agglutination assay kit, (ID-Anti-Fya/Fyb, Biorad), was utilized to assess the Duffy phenotype.

### *In vitro* invasion inhibition assay using *P. vivax* clinical isolates

Cryopreserved *P. vivax* isolates collected from Cambodian patients infected *P. vivax* were used for *in vitro* invasion inhibition assays (54) as described in Supplementary Text S4. The mouse monoclonal anti-Duffy antibody 2C3 and pre-immune rabbit sera were used as positive and negative controls, respectively. After incubation, cells were stained with Hoechst 33342 to label parasite DNA and parasitemia was quantified by flow cytometry (BD FacsAria Fusion™). Reticulocytes that were positive for both Hoechst 33342 and Cell Trace Far Red were scored as newly invaded cells. For analysis, data were normalized to invasion levels observed in the absence of antibodies and analyzed using FlowJo (v10.8.1) software. Invasion rates in the absence of antibodies ranged from 0.3% to 3.9%. Use of *P. vivax* clinical isolates for *ex vivo* invasion assays and preparation of IFA slides was approved by the Cambodian National Ethics Committee for Health Research (# 317-NECHR).

### Immunoprecipitation of blood stage antigens and detection of interacting proteins by mass spectrometry

*P. knowlesi* merozoites were isolated from synchronized blood stage cultures as described above. Merozoite surface proteins were crosslinked by treatment with 2mM dithiobis succinimidyl propionate (DSP) at room temperature for 30 mins . Merozoite lysates were prepared by adding lysis buffer (50 mM Tris, 150 mM NaCl, 1 mM EDTA, 1% NP-40, 5% glycerol (pH 7.4)), incubating on ice for 1h, and mechanically lysing the cells by passing through a 26-gauge needle. Finally, the lysate was centrifuged at 19,500 g for 40 minutes at 4℃. *P. knowlesi* merozoite lysate was pre-cleared using protein G-magnetic beads prior to incubation with protein G-magnetic beads (40 µl) loaded with anti-PvMP45, anti-PvMP36, or pre-immune rabbit sera. Protein G-magnetic beads were then washed with buffer containing 0.1% Tween and split into two halves, with the first half used to elute bound proteins for western blot analysis. The other half was washed three times with 0.1 M ammonium bicarbonate, treated with 0.2 μg of trypsin-LysC (Promega) for 1 h at 37 °C. Resulting peptides were desalted, lyophilized and reconstituted in 10 µl of injection buffer in 0.3 % trifluoroacetic acid for analysis by liquid chromatography-tandem mass spectrometry (LC-MS/MS) as described in Suppplementary Text S5. Data from LC-MS/MS analysis was analyzed to identify interacting proteins as described in Supplementary Text S6. The mass spectrometry proteomics raw data have been deposited in the ProteomeXchange Consortium via the PRIDE partner repository (55,56) with the dataset identifier PXD075937. Initially, the two newly identified genes were designated as BPV85 and BPV86. Later, based on their microneme localization and molecular weight, they were renamed PvMP45 and PvMP36 respectively. In the MyProMS software analysis, immunoprecipitation experiments using PvMP45 and PvMP36 identified 202 and 134 proteins, respectively. Proteins were selected based on a fold change > 1.3, an adjusted P-value > 0.05, and the identification of at least two peptides across three replicates. Proteins included in Tables 1 and 2 were further selected based on known or predicted role in erythrocyte invasion or known localization on the merozoite surface or in invasion related apical organelles such as micronemes and rhoptries.

### Protein-Protein interaction studies using ELISA based binding assay

Binding interactions between recombinant PvMP45 and PvMP36 with PkRAP1 were tested as described below. Recombinant PvMP45 and PvMP36 proteins in 100 mM carbonate/bicarbonate buffer were coated on ELISA wells (100 ng/well). Recombinant PkRAP1 was incubated at different concentrations (0-5 µg/ml) in wells coated with PvMP45 and PvMP36. Unbound protein was washed away and bound protein was detected using specific antisera in an ELISA format. Recombinant PvDBPII was used as negative control. All experiments were performed in duplicate with three biological repeats, and the means ± standard error of mean (SEM) are reported.

## ACKNOWLEDGEMENTS

We sincerely appreciate all individuals and their families living in malaria endemic regions of Papua New Guinea and Thailand who participated in this study. We also extend our gratitude to the extensive teams in Papua New Guinea and Thailand for their dedication and effort in conducting the field studies. CEC was supported by ANR Grants, PvINV (ANR-21-CE15-0013-01) and Labex PRAFRAP (ANR-11-LABX-0024-PARAFRAP). RJL was supported by a NHMRC Investigator Grant (GNT1173210), the Victorian Government as a veski FAIR Fellow and from The Sylvia and Charles Viertel Charitable Foundation as a Viertel Senior Medical Research Fellow. The authors acknowledge the Victorian State Government Operational Infrastructure Support and Australian Government NHMRC IRIISS. JP was supported by NIH (R01AI173171, R01AI175134 and R01AI143694) and the Pasteur International Unit PvESMEE.

## AUTHOR CONTRIBUTIONS

AD and FM designed and performed experiments, analyzed data, and wrote the manuscript. PSL, LBFD, KP, BT, BK, DHO performed experiments and analyzed data. YLL, MYF, ETH provided valuable reagents. JGB, JS, IM and JP supervised research and analyzed data. RL designed field studies, supervised research and analyzed data. CEC designed and co-ordinated the study, supervised research, obtained funding and wrote the manuscript. All authors reviewed and contributed to the manuscript and approved the final version.

## REFERENCES

1. World Health Organization. World malaria report 2025 (World Health Organization, 2025). https://www.who.int/teams/global-malaria-programme/reports/world-malaria-report-2025

2. Miller LH, Mason SJ, Clyde DF, McGinniss MH. 1976. The resistance factor to *Plasmodium vivax* in blacks: the Duffy-blood-group genotype, FyFy. N Engl J Med 295:302–304. 10.1056/NEJM197608052950602

3. Ménard D, Barnadas C, Bouchier C, Henry-Halldin C, Gray LR, Ratsimbasoa A, Thonier V, Carod J, Domarle O, Colin Y, Bertrand O, Picot J, King CL, Grimberg BT, Mercereau-Puijalon O, Zimmerman PA.2010. *Plasmodium vivax* clinical malaria is cmmonly observed in Duffy- negative Malagasy people. Proc Natl Acad Sci U S A 107:5967–5971. 10.1073/pnas.0912496107

4. Ngassa Mbenda HG, Das A. 2014. Molecular evidence of *Plasmodium vivax* mono and mixed infections in Duffy-negative native Cameroonians. PLoS One 9:e103262. 10.1371/journal.pone.0103262

5. Woldearegai TG, Kremsner PG, Kun JF, Mordmüller B. 2013. *Plasmodium vivax* malaria in Duffy-negative individuals from Ethiopia. Trans R Soc Trop Med Hyg 107:328–331. 10.1093/trstmh/trt016

6. Abdelraheem MH, Albsheer MM, Mohamed HS, Amin M, Abdel Hamid MM. 2016. Transmission of *Plasmodium vivax* in Duffy-negative individuals in central Sudan. Trans R Soc Trop Med Hyg 110:258–260. 10.1093/trstmh/trw014

7. Bouyssou I, El Hoss S, Doderer-Lang C, Schoenhals M, Rasoloharimanana LT, Vigan-Womas I, Ratsimbasoa A, Abate A, Golassa L, Mabilotte S, Kessler P, Guillotte-Blisnick M, Martinez FJ, Chitnis CE, Strouboulis J, Ménard D. 2023. Unveiling *P. vivax* invasion pathways in Duffy- negative individuals. Cell Host Microbe. 31:2080–2092.e5. 10.1016/j.chom.2023.11.007

8. Bouyssou I, El Hoss S, Doderer-Lang C, Schoenhals M, Rasoloharimanana LT, Vigan-Womas I, Ratsimbasoa A, Rees DC, Abate A, Golassa L, Mabilotte S, Guillotte M, Martinez Blazquez FJ, Chitnis CE, Strouboulis J, Ménard D. 2023. The Darc side of *Vivax* malaria in Africa: unveiling invasion pathways into Duffy-negative erythroblasts. mBio 14(1):e184938. 10.1182/blood-2023-184938

9. Dechavanne C, Dechavanne S, Bosch J, Metral S, Redinger KR, Watson QD, Ratsimbasoa AC, Roeper B, Krishnan S, Fong R, Bennett S, Carias L, Chen E, Salinas ND, Ghosh A, Tolia NH, Woost PG, Jacobberger JW, Colin Y, Gamain B, King CL, Zimmerman PA. 2023. Duffy antigen is expressed during erythropoiesis in Duffy-negative individuals. Cell Host Microbe 31:2093–2106.e7. 10.1016/j.chom.2023.10.019

10. Battle KE, Baird JK. 2021. The global burden of *Plasmodium vivax* malaria is obscure and insidious. PLoS Med. 2021;18(12):e1003799. 10.1371/journal.pmed.1003799

11. Osoro CB, Ochodo E, Kwambai TK, Otieno JA, Were L, Sagam CK, Owino EJ, Kariuki S, Ter Kuile FO, Hill J. 2024. Policy uptake and implementation of the RTS,S/AS01 malaria vaccine in sub-Saharan African countries: status 2 years following the WHO recommendation. BMJ Glob Health 9:e014719. 10.1136/bmjgh-2023-014719

12. von Seidlein L. 2025. Roll out and prospects of the malaria vaccine R21/Matrix-M. PLoS Med 22:e1004515. 10.1371/journal.pmed.1004515

13. Salman AM, Montoya-Díaz E, West H, Lall A, Atcheson E, Lopez-Camacho C, Ramesar J, Bauza K, Collins KA, Brod F, Reis F, Pappas L, González-Cerón L, Janse CJ, Hill AVS, Khan SM, Reyes-Sandoval A. 2017. Rational development of a protective *Plasmodium vivax* vaccine evaluated with transgenic rodent parasite challenge models. Sci Rep 7:46482. 10.1038/srep46482

14. Hou MM, Barrett JR, Themistocleous Y, Rawlinson TA, Diouf A, Martinez FJ, Nielsen CM, Lias AM, King LDW, Edwards NJ, Greenwood NM, Kingham L, Poulton ID, Khozoee B, Goh C, Hodgson SH, Mac Lochlainn DJ, Salkeld J, Guillotte-Blisnick M, Huon C, Mohring F, Reimer JM, Chauhan VS, Mukherjee P, Biswas S, Taylor IJ, Lawrie AM, Cho J-S, Nugent FL, Long CA, Moon RW, Miura K, Silk SE, Chitnis CE, Minassian AM, Draper SJ. 2023. Vaccination with *Plasmodium vivax* Duffy-binding protein inhibits parasite growth during controlled human malaria infection. Sci Transl Med 15:eadf1782. 10.1126/scitranslmed.adf1782

15. Martinez FJ, White M, Guillotte-Blisnick M, Huon C, Boucharlat A, Agou F, England P, Popovici J, Hou MM, Silk SE, Barrett JR, Nielsen CM, Reimer JM, Mukherjee P, Chauhan VS, Minassian AM, Draper SJ, Chitnis CE. 2024. PvDBPII elicits multiple antibody-mediated mechanisms that reduce growth in a *Plasmodium vivax* challenge trial. NPJ Vaccines 9:10. 10.1101/2023.08.01.23293515

16. Hester J, Chan ER, Menard D, Mercereau-Puijalon O, Barnwell J, Zimmerman PA, Serre D. 2013. De novo assembly of a field isolate genome reveals novel Plasmodium vivax erythrocyte invasion genes. PLoS Negl Trop Dis. 7(12):e2569. 10.1371/journal.pntd.0002569

17. Roesch C, Popovici J, Bin S, Run V, Kim S, Ramboarina S, Rakotomalala E, Rakotoarison RL, Rasoloharimanana T, Andriamanantena Z, Kumar A, Guillotte-Blisnick M, Huon C, Serre D, Chitnis CE, Vigan-Womas I, Menard D. 2018. Genetic diversity in two *Plasmodium vivax* protein ligands for reticulocyte invasion. PLoS Negl Trop Dis 12:e0006555. 10.1371/journal.pntd.0006555

18. Han JH, Cho JS, Ong JJY, Park JH, Nyunt MH, Sutanto E, Trimarsanto H, Petros B, Aseffa A, Getachew S, Sriprawat K, Anstey NM, Grigg MJ, Barber BE, William T, Qi G, Liu Y, Pearson RD, Auburn S, Price RN, Nosten F, Rénia L, Russell B, Han ET. 2020. Genetic diversity and neutral selection in *Plasmodium vivax* erythrocyte binding protein correlates with patient antigenicity. mBio 11(3):e0008202. 10.1371/journal.pntd.0008202

19. Lee SK, Crosnier C, Valenzuela-Leon PC, Dizon BLP, Atkinson JP, Mu J, Wright GJ, Calvo E, Gunalan K, Miller LH. 2024. Complement receptor 1 is the human erythrocyte receptor for *Plasmodium vivax* erythrocyte binding protein. Proc Natl Acad Sci U S A 121:e2316304121. 10.1073/pnas.2316304121

20. King CL, Michon P, Shakri AR, Marcotty A, Stanisic D, Zimmerman PA, Cole-Tobian JL, Mueller I, Chitnis CE. 2008. Naturally acquired Duffy-binding protein-specific binding inhibitory antibodies confer protection from blood-stage Plasmodium vivax infection. Proc Natl Acad Sci U S A. 105(24):8363–8. doi: 10.1073/pnas.0800371105. https://doi.org/10.1073/pnas.0800371105

21. He WQ, Shakri AR, Bhardwaj R, França CT, Stanisic DI, Healer J, Kiniboro B, Robinson LJ, Guillotte-Blisnick M, Huon C, Siba P, Cowman A, King CL, Tham WH, Chitnis CE, Mueller I. 2019. Antibody responses to *Plasmodium vivax* Duffy binding and Erythrocyte binding proteins predict risk of infection and are associated with protection from clinical Malaria. PLoS Negl Trop Dis. 13(2):e0006987. 10.1371/journal.pntd.0006987

22. França CT, White MT, He WQ, Hostetler JB, Brewster J, Frato G, Malhotra I, Gruszczyk J, Huon C, Lin E, Kiniboro B, Yadava A, Siba P, Galinski MR, Healer J, Chitnis C, Cowman AF, Takashima E, Tsuboi T, Tham WH, Fairhurst RM, Rayner JC, King CL, Mueller I. 2017. Identification of highly-protective combinations of *Plasmodium vivax* recombinant proteins for vaccine development. Elife. 6:e28673. 10.7554/eLife.28673

23. Crosnier C, Bustamante LY, Bartholdson SJ, Bei AK, Theron M, Uchikawa M, Mboup S, Ndir O, Kwiatkowski DP, Duraisingh MT, Rayner JC, Wright GJ. 2011. Basigin is a receptor essential for erythrocyte invasion by *Plasmodium falciparum*. Nature 480:534–537. 10.1038/nature10606

24. Wright KE, Hjerrild KA, Bartlett J, Douglas AD, Jin J, Brown RE, Illingworth JJ, Ashfield R, Clemmensen SB, de Jongh WA, Draper SJ, Higgins MK. 2014. Structure of malaria invasion protein RH5 with erythrocyte basigin and blocking antibodies. Nature 515:427–430. 10.1038/nature13715

25. Chen L, Xu Y, Wong W, Thompson JK, Healer J, Goddard-Borger ED, Lawrence MC, Cowman AF. 2017. Structural basis for inhibition of erythrocyte invasion by antibodies to *Plasmodium falciparum* protein CyRPA. eLife 6:e21347. 10.7554/eLife.21347

26. Volz JC, Yap A, Sisquella X, Thompson JK, Lim NTY, Whitehead LW, Chen L, Lampe M, Tham WH, Wilson D, Nebl T, Marapana D, Triglia T, Wong W, Rogers KL, Cowman AF. 2016. Essential role of the PfRh5/PfRipr/CyRPA complex during *Plasmodium falciparum* invasion of erythrocytes. Cell Host Microbe 20:60–71. 10.1016/j.chom.2016.06.004

27. Farrell B, Alam N, Hart MN, Jamwal A, Ragotte RJ, Walters-Morgan H, Draper SJ, Knuepfer E, Higgins MK. 2024. The PfRCR complex bridges malaria parasite and erythrocyte during invasion. Nature 625:578–584. 10.1038/s41586-023-06856-1

28. Scally SW, Triglia T, Evelyn C, Seager BA, Pasternak M, Lim PS, Healer J, Geoghegan ND, Adair A, Tham WH, Dagley LF, Rogers KL, Cowman AF. 2022. PCRCR complex is essential for invasion of human erythrocytes by *Plasmodium falciparum*. Nat Microbiol 7:2039–2053. 10.1038/s41564-022-01261-2

29. Knuepfer E, Wright KE, Prajapati SK, Rawlinson TA, Mohring F, Koch M, Lyth OR, Howell SA, Villasis E, Snijders AP, Moon RW, Draper SJ, Rosanas-Urgell A, Higgins MK, Baum J, Holder AA. 2019. Divergent roles for the RH5 complex components, CyRPA and RIPR, in human- infective malaria parasites. PLoS Pathog 15:e1007809. 10.1371/journal.ppat.1007809

30. Seager BA, Lim PS, Xiao X, Lai KH, Feufack-Donfack LB, Dass S, Jung NC, Abraham A, Grigg MJ, Anstey NM, William T, Sattabongkot J, Leis A, Longley RJ, Duraisingh MT, Popovici J, Wilson DW, Cowman AF, Scally SW. 2026. PTRAMP, CSS and Ripr form a conserved complex required for merozoite invasion of *Plasmodium* species into erythrocytes. Nature Communications 17:1780. 10.1038/s41467-026-68486-1

31. Elsworth B, Ye S, Dass S, Tennessen JA, Sultana Q, Thommen BT, Paul AS, Kanjee U, Grüring C, Ferreira MU, Gubbels MJ, Zarringhalam K, Duraisingh MT. 2025. The essential genome of *Plasmodium knowlesi* reveals determinants of antimalarial susceptibility. Science 387:eadq6241. 10.1126/science.adq6241

32. Zhang M, Wang C, Otto TD, Oberstaller J, Liao X, Adapa SR, Udenze K, Bronner IF, Casandra D, Mayho M, Brown J, Li S, Swanson J, Rayner JC, Jiang RHY, Adams JH. 2018. Uncovering the essential genes of the human malaria parasite *Plasmodium falciparum* by saturation mutagenesis. Science 360:eaap7847. 10.1126/science.aap7847

33. Aurrecoechea C, Brestelli J, Brunk BP, Dommer J, Fischer S, Gajria B, Gao X, Gingle A, Grant G, Harb OS, Heiges M, Innamorato F, Iodice J, Kissinger JC, Kraemer E, Li W, Miller JA, Nayak V, Pennington C, Pinney DF, Roos DS, Ross C, Stoeckert CJ Jr, Treatman C, Wang H. 2009. PlasmoDB: a functional genomic database for malaria parasites. Nucleic Acids Res 37:D539–D543. 10.1093/nar/gkn814

34. Krogh A, Larsson B, von Heijne G, Sonnhammer EL. 2001. Predicting transmembrane protein topology with a hidden Markov model: application to complete genomes. J Mol Biol 305:567–580. 10.1006/jmbi.2000.4315

35. Hallgren J, Tsirigos KD, Pedersen MD, Almagro Armenteros JJ, Marcatili P, Nielsen H, Krogh A, Winther O. 2022. DeepTMHMM predicts alpha and beta transmembrane proteins using deep neural networks. bioRxiv. 10.1101/2022.04.08.487609

36. Schultz J, Copley RR, Doerks T, Ponting CP, Bork P. 2000. SMART: a web-based tool for the study of genetically mobile domains. Nucleic Acids Res 28:231–234. 10.1093/nar/28.1.231

37. Kelley LA, Mezulis S, Yates CM, Wass MN, Sternberg MJE. 2015. The Phyre2 web portal for protein modeling, prediction and analysis. Nat Protoc 10:845–858. 10.1038/nprot.2015.053

38. Wagner MP, Formaglio P, Gorgette O, Dziekan JM, Huon C, Berneburg I, Rahlfs S, Barale JC, Feinstein SI, Fisher AB, Ménard D, Bozdech Z, Amino R, Touqui L, Chitnis CE. 2022. Human peroxiredoxin 6 is essential for malaria parasites and provides a host-based drug target. Cell Rep. 39(11):110923. 10.1016/j.celrep.2022.110923

39. Deshmukh A, Chourasia BK, Mehrotra S, Kana IH, Paul G, Panda A, Kaur I, Singh SK, Rathore S, Das A, Gupta P, Kalamuddin M, Gakhar SK, Mohmmed A, Theisen M, Malhotra P. 2018. *Plasmodium falciparum* MSP3 exists in a complex on the merozoite surface and generates antibody response during natural infection. Infect Immun 86:e00067–18. 10.1128/iai.00067-18

40. Hostetler JB, Sharma S, Bartholdson SJ, Wright GJ, Fairhurst RM, Rayner JC. 2015. A library of *Plasmodium vivax* recombinant merozoite proteins reveals new vaccine candidates and protein–protein interactions. PLoS Negl Trop Dis 9:e0004264. 10.1371/journal.pntd.0004264

41. Osier FH, Feng G, Boyle MJ, Langer C, Zhou J, Richards JS, McCallum FJ, Reiling L, Jaworowski A, Anders RF, Marsh K, Beeson JG. 2014. Opsonic phagocytosis of *Plasmodium falciparum* merozoites: mechanism in human immunity and a correlate of protection against malaria. BMC Med 12:108. 10.1186/1741-7015-12-108

42. Opi DH, Longley RJ, Takashima E, Spelman T, Tayipto Y, Schoffer K, Brewster J, Reiling L, Wines BD, Kiniboro B, Siba P, Harbers M, Hogarth M, Tsuboi T, Robinson LJ, Mueller I, Beeson JG. 2026. A longitudinal study of children identified antibody Fc-mediated functions and antigen targets of immunity to *Plasmodium vivax* malaria. Immunity. S1074-7613(26)00070-1. 10.1016/j.immuni.2026.02.003

43. Nkumama IN, Ogwang R, Odera D, Musasia F, Mwai K, Nyamako L, Murungi L, Tuju J, Fürle K, Rosenkranz M, Kimathi R, Njuguna P, Hamaluba M, Kapulu MC, Frank R, CHMI-SIKA Study Team, Osier FHA. 2024. Breadth of Fc-mediated effector function correlates with clinical immunity following human malaria challenge. Immunity 57:1215–1224.e6. 10.1016/j.immuni.2024.05.001.

44. McIntosh RS, Shi J, Jennings RM, Chappel JC, de Koning-Ward TF, Smith T, Green J, van Egmond M, Leusen JHW, Lazarou M, van de Winkel JGJ, Jones TS, Crabb BS, Holder AA, Pleass RJ. 2007. The importance of human FcγRI in mediating protection to malaria. PLoS Pathog 3:e72. 10.1371/journal.ppat.0030072

45. Moon RW, Hall J, Rangkuti F, Ho YS, Almond N, Mitchell GH, Pain A, Holder AA, Blackman MJ. 2013. Adaptation of the genetically tractable malaria pathogen *Plasmodium knowlesi* to continuous culture in human erythrocytes. Proc Natl Acad Sci U S A 110:531–536.

46. Lyth O, Vizcay-Barrena G, Wright KE, Haase S, Mohring F, Najer A, Henshall IG, Ashdown GW, Bannister LH, Drew DR, Beeson JG, Fleck RA, Moon RW, Wilson DW, Baum J. 2018. Cellular dissection of malaria parasite invasion of human erythrocytes using viable *Plasmodium knowlesi* merozoites. Sci Rep 8:10165. 10.1038/s41598-018-28457-z.

47. Lin E, Kiniboro B, Gray L, Dobbie S, Robinson L, Laumaea A, Schöpflin S, Stanisic D, Betuela I, Blood-Zikursh M, Siba P, Felger I, Schofield L, Zimmerman P, Mueller I. 2010. Differential patterns of infection and disease with P. falciparum and P. vivax in young Papua New Guinean children. PLoS One. 5(2):e9047. 10.1371/journal.pone.0009047

48. Longley RJ, Sripoorote P, Chobson P, Saeseu T, Sukasem C, Phuanukoonnon S, Nguitragool W, Mueller I, Sattabongkot J. 2016. High Efficacy of Primaquine Treatment for Plasmodium vivax in Western Thailand. Am J Trop Med Hyg. 95(5):1086–1089. 10.4269/ajtmh.16-0410

49. Mazhari R, Brewster J, Fong R, Bourke C, Liu ZSJ, Takashima E, Tsuboi T, Tham WH, Harbers M, Chitnis C, Healer J, Ome-Kaius M, Sattabongkot J, Kazura J, Robinson LJ, King C, Mueller I, Longley RJ. 2020. A comparison of non-magnetic and magnetic beads for measuring IgG antibodies against *Plasmodium vivax* antigens in a multiplexed bead-based assay using Luminex technology (Bio-Plex 200 or MAGPIX). PLoS One 15:e0238010. 10.1371/journal.pone.0238010

50. Liu ZSJ, Sattabongkot J, White M, Chotirat S, Kumpitak C, Takashima E, Harbers M, Tham WH, Healer J, Chitnis CE, Tsuboi T, Mueller I, Longley RJ. 2022. Naturally acquired antibody kinetics against *Plasmodium vivax* antigens in people from a low malaria transmission region in western Thailand. BMC Med 20:89. 10.1186/s12916-022-02281-9

51. Pekin K. 2025. Opsonic phagocytosis assay using the THP-1 monocytic cell line and antigen- coated beads. 10.17504/protocols.io.ewov19pwylr2/v1

52. Singh S, Pandey K, Chattopadhyay R, Yazdani SS, Lynn A, Bharadwaj A, Ranjan A, Chitnis CE. 2001. Biochemical, biophysical, and functional characterization of bacterially expressed and refolded receptor binding domain of *Plasmodium vivax* Duffy-binding protein. J Biol Chem 276:17111–17116. 10.1074/jbc.m101531200

53. Yazdani SS, Shakri AR, Mukherjee P, Baniwal SK, Chitnis CE. 2004. Evaluation of immune responses elicited in mice against a recombinant malaria vaccine based on *Plasmodium vivax* Duffy binding protein. Vaccine 22:3727–3737. 10.1016/j.vaccine.2004.03.030

54. Popovici J, Roesch C, Carias LL, Khim N, Kim S, Vantaux A, Mueller I, Chitnis CE, King CL, Witkowski B. 2020. Amplification of Duffy binding protein-encoding gene allows Plasmodium vivax to evade host anti-DBP humoral immunity. Nat Commun. 11(1):953. 10.1038/s41467-020-14574-9

55. Perez-Riverol Y. 2022. Proteomic repository data submission, dissemination, and reuse: key messages. Expert Rev Proteomics 19:297–310. 10.1080/14789450.2022.2160324

56. Poullet P, Carpentier S, Barillot E. 2007. myProMS, a web server for management and validation of mass spectrometry-based proteomic data. Proteomics 7:2553–2556. 10.1002/pmic.200600784

